# Individual-Based Network Model for Rift Valley Fever in Kabale District, Uganda, to Guide Mitigation Measures: A One Health Model

**DOI:** 10.1101/388785

**Authors:** Musa Sekamatte, Mahbubul H. Riad, Tesfaalem Tekleghiorghis, Kenneth J. Linthicum, Seth C. Britch, Juergen A. Richt, J. P. Gonzalez, Caterina M Scoglio

## Abstract

Rift Valley fever (RVF) is a zoonotic disease which causes significant morbidity and mortality among ungulate livestock and humans in endemic regions. In the major RVF epizootic regions of East Africa, the causative agent of the disease, Rift Valley fever virus (RVFV), is primarily transmitted by multiple mosquito species in *Aedes, Culex*, and *Mansonia* genera during both epizootic and enzootic periods in a complex transmission cycle largely driven by the environment. However, recent RVFV activity in Uganda demonstrated that RVFV could also spread into new regions through livestock movements, and underscored the need to develop effective mitigation strategies to reduce transmission and prevent spread among cattle operations. We simulated RVFV transmission among cattle in different sub counties of Kabale District in Uganda using real world livestock data in a network-based model. This model considered livestock as spatially explicit factors in different sub-counties subjected to specific vector mosquito and environmental factors, and was configured to investigate and quantitatively evaluate the relative impacts of mosquito control, livestock movement regulations, and diversity in cattle populations on the spread of the RVF epizootic. We concluded that cattle movement should be restricted during periods of high vector mosquito abundance to control the epizootic spreading among sub-counties. On the other hand we found that mosquito control would only be sufficient to control the epizootic when mosquito abundance was low. Importantly, simulation results also showed that cattle populations with a higher diversity with regard to indigenous combined with exotic breeds led to reduced numbers of infected cattle compared to more homogenous cattle populations.

## Introduction

Rift Valley fever (RVF) is a zoonotic mosquito-borne disease caused by Rift Valley fever virus (RVFV; *Phlebovirus:* Bunyaviridae) that severely affects ungulate livestock and wildlife but can also affect humans in RVF-endemic regions of sub-Saharan Africa and parts of the Arabian Peninsula [1, 2]. The potential extreme economic, and public and veterinary health burdens of epizootics/epidemics of RVF have been described extensively [3]. Heavy rainfall and flooding are the most prominent precursors of RVF epizootics in savanna areas of Africa, due to flooded ground pools stimulating massive emergence of transovarially RVFV-infected *Aedes* mosquitoes and the initiation of the transmission cycle as the virus is introduced into livestock by these mosquitoes during blood feeding [2, 4, 5]. However, areas outside the recognized RVF epizootic regions, especially in central Africa, may not experience transmissions linked to elevated rainfall [6]. In these areas, RVFV is most likely spread via movements of infected livestock from endemic areas with elevated rainfall: livestock trading across different market areas may include infectious cattle that could disperse the virus and, in the presence of suitable mosquito vectors, accelerate and expand the transmission of RVFV, especially when cattle operations are distributed across large distances but linked by trade [7]. Patterns of recent RVF activity in Uganda first described on 9 March 2016 support this hypothesis of RVFV spread linked to the cattle trade [8] and underscore the need to develop effective operational surveillance and mitigation strategies to reduce transmission and prevent spread among cattle operations. In this study we designed a network-based epidemic transmission model to run simulations to quantitatively investigate the patterns of spread of RVFV across cattle operations in Kabale District, Uganda, providing an opportunity to thus quantitatively investigate the potential impact of various mitigation methods.

Scoglio et al. [9] recently published an individual-level RVF epidemic transmission model for cattle in Riley County, Kansas, USA, structured on the *susceptible-exposed-infected-recovered* (SEIR) framework applied to a cattle movement network. They used two separate kernel functions – exponential and power-law models – to model cattle movement between and among farms in Riley County. Simulations with these kernel functions revealed that more widespread epizootics resulted from the power law model, most likely because cattle were allowed to move to distant farms. In contrast, the exponential model greatly restricted cattle movement to more proximal farms, reducing spread of the virus. The message from the Scoglio et al. study is that restricting cattle movement substantially reduces RVFV transmission and spread across the landscape of cattle operations. Secondarily, they found that partitioning each farm into several clusters also results in less widespread RVF epizootics.

In the present study, we built upon the Scoglio et al. [9] model to investigate RVFV epidemiology in Kabale District using real world livestock data. This model considered livestock as spatially explicit factors in an individual-based network representing different sub-counties subjected to specific mosquito and environmental factors. We designed the cattle movement network based on the local trading system for the Kabale District, but considered two different networks depending upon the relative susceptibility of exotic and indigenous cattle.

Mosquito vectors for RVFV were included in an aggregated way with a parameter for transmission rate from an infectious animal to a susceptible one, which included mosquito abundance, survival rate, vector competence, and feeding patterns – collectively proportional to vectorial capacity, the efficiency of vector borne disease transmission [9]. Simulations were performed for a variety of initial outbreak conditions, such as single location versus multi-location initial outbreaks, as well as outbreaks in locations with varying cattle populations, mosquito transmission rates, and different cattle movement probabilities among sub-counties. We configured the model to investigate and quantitatively evaluate the relative impacts of mosquito control, livestock movement regulations, and diversity – that is, whether cattle are indigenous or exotic breeds – in cattle populations on containing the spread of the RVF epizootic. Materials and Methods

### Modeling Framework

A simplified diagram of the individual-based network modeling framework we used to model RVFV epidemiology in Kabale District, Uganda, is presented in **Fig. 1**. Our RVFV modeling framework consists of two parts, a node transition graph and a contact network. The node transition graph consists of four compartments— Susceptible (*S*), Exposed (*E*), Infectious (*I*), and Recovered (*R*). Once a susceptible cow gets bitten by a RVFV-infected mosquito (exposed), there is a possibility that after a certain period of viral replication, the cow itself becomes infectious. The infectious cow can in turn infect susceptible naïve mosquitoes who bite it during the period before the cow recovers or is removed, for example by death or culling. Each individual cow can be in only one of these four compartments and rates of transitions between compartments are driven by parameters *β* (transmission rate), *δ* (extrinsic incubation period of the virus), and *γ* (recovery rate) in **Fig. 1**. The contact network consists of the total population size (*N*) of cows in the network, each represented by a small square, or node, and a black line links two nodes when an opportunity for transmission of RVFV between two cows (via the bite of an infectious mosquito) arises.

**Fig. 1:**
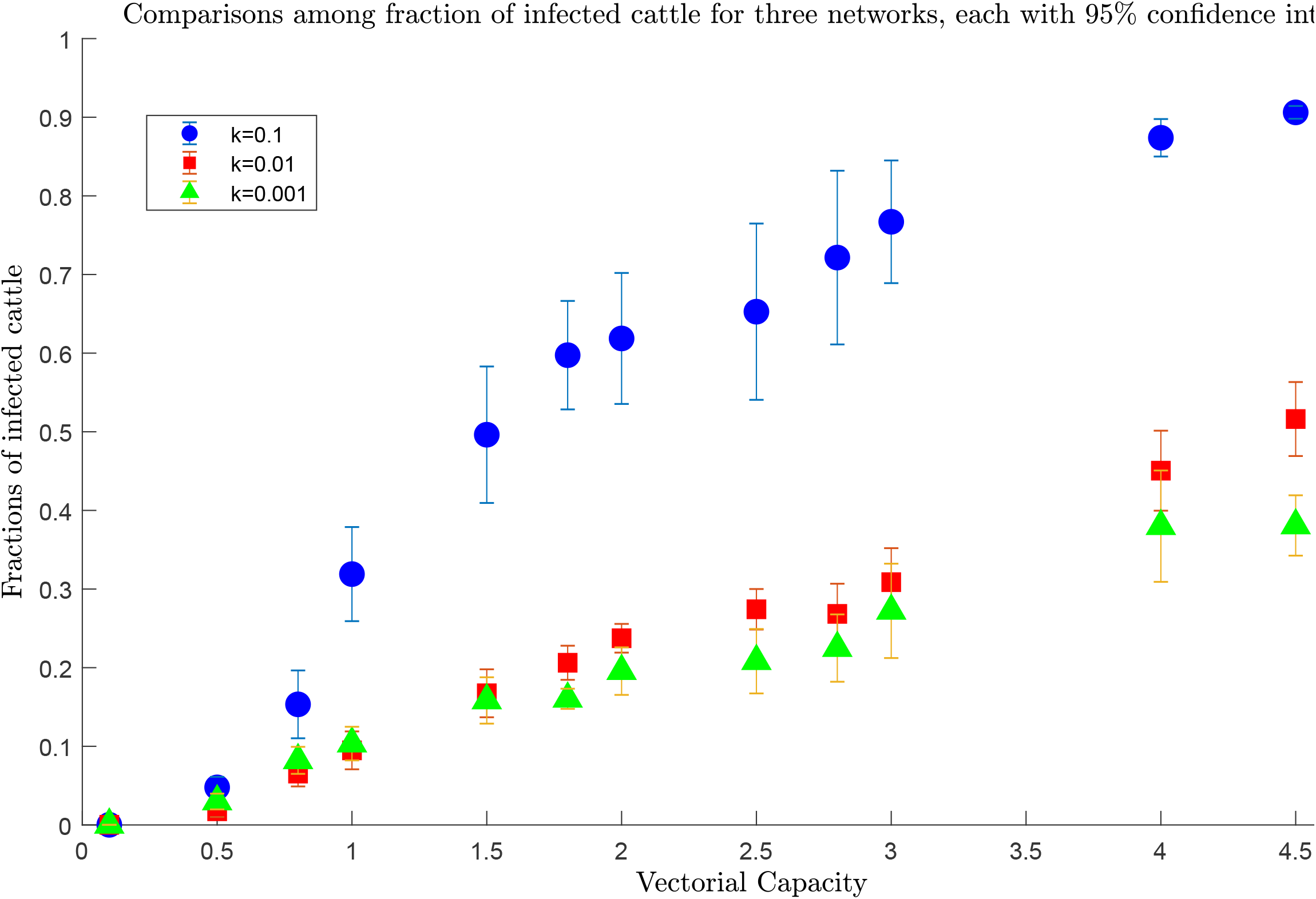
Simplified diagram of an individual-based network model which consists of a node transition graph and a contact network. **Circles in the node transition graph represent the four compartments** Susceptible (*S*), Exposed (*E*), Infectious (*I*), and Recovered (*R*) of a node (i.e., of an individual cow), and **arrows between the compartments show the direction of transition for each node (cow)** with rates driven by parameters *β* (transmission rate), *δ* (extrinsic incubation period of the virus), and *γ* (recovery rate). **Circles in the contact network in turn represent individual cows (i.e., nodes), and black lines linking circles represent opportunities for RVFV transmission**.

### Geographic Structure and Movement in the Cattle Contact Network

The cattle contact network consisted of 20,806 cattle (N) unevenly distributed across 19 sub-counties in the Kabale District of Uganda (**Table 1** and **Fig. 2**), which is approximately 1,679 km^2^ (648 sq mi) with a human population of nearly half a million in the western region of Uganda. We obtained these cattle abundance data, as well as the proportions of indigenous and exotic cows in each sub-district. From the Uganda Bureau of Statistics (UBOS) [10].

**Table I:**
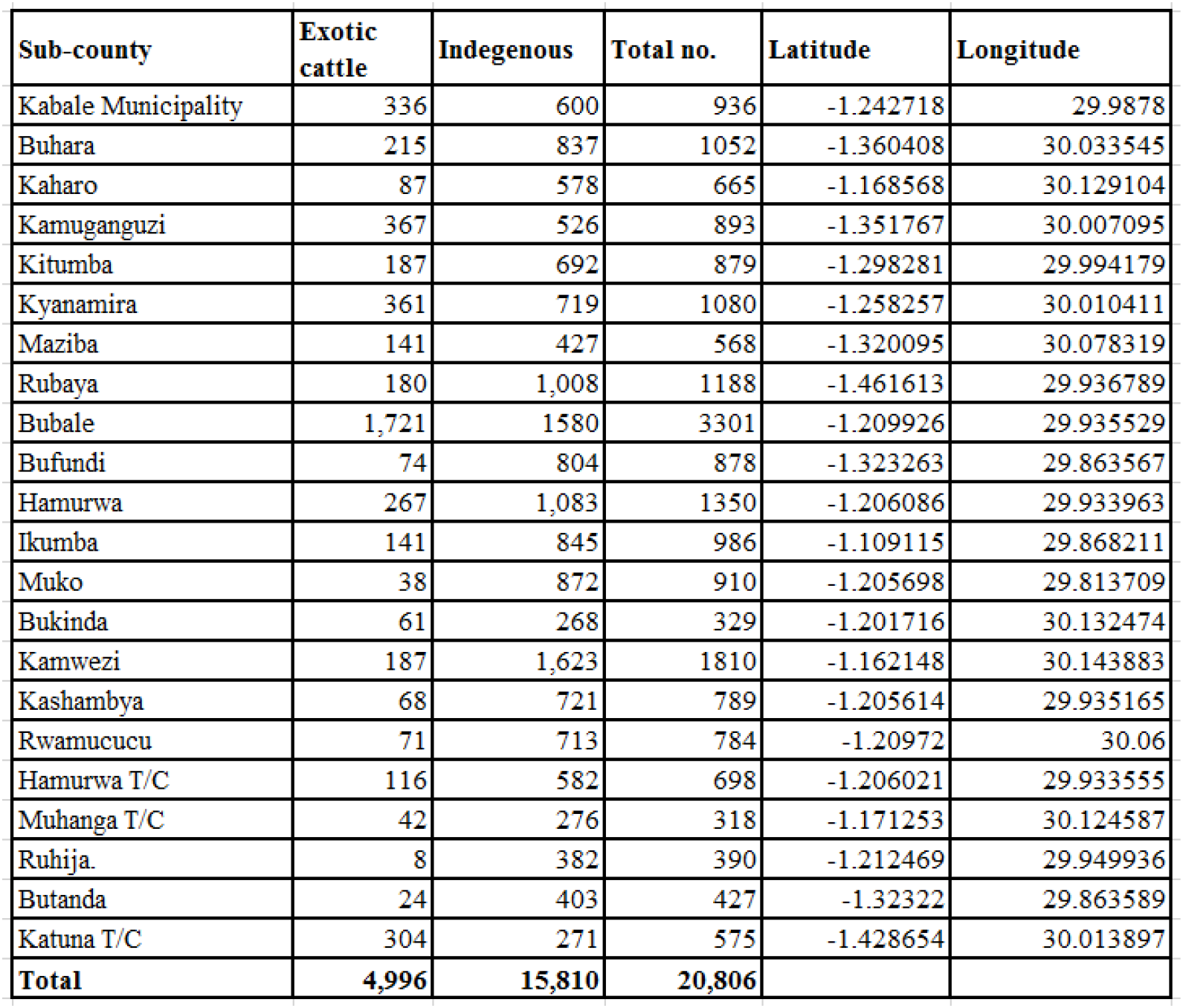
Cattle in different sub-counties in Kabale District. This data set was derived from UBOS Statistical Report 012, Kabale District [10]

| Sub-county | Exotic cattle | Indigenous | Total no. | Latitude | Longitude |
| --- | --- | --- | --- | --- | --- |
| Kabale Municipality | 336 | 600 | 936 | -1.242718 | 29.9878 |
| Buhara | 215 | 837 | 1052 | -1.360408 | 30.033545 |
| Kaharo | 87 | 578 | 665 | -1.168568 | 30.129104 |
| Kamuganguzi | 367 | 526 | 893 | -1.351767 | 30.007095 |
| Kitumba | 187 | 692 | 879 | -1.298281 | 29.994179 |
| Kyanamira | 361 | 719 | 1080 | -1.258257 | 30.010411 |
| Maziba | 141 | 427 | 568 | -1.320095 | 30.078319 |
| Rubaya | 180 | 1,008 | 1188 | -1.461613 | 29.936789 |
| Bubale | 1,721 | 1580 | 3301 | -1.209926 | 29.935529 |
| Bufundi | 74 | 804 | 878 | -1.323263 | 29.863567 |
| Hamurwa | 267 | 1,083 | 1350 | -1.206086 | 29.933963 |
| Ikumba | 141 | 845 | 986 | -1.109115 | 29.868211 |
| Muko | 38 | 872 | 910 | -1.205698 | 29.813709 |
| Bukinda | 61 | 268 | 329 | -1.201716 | 30.132474 |
| Kamwezi | 187 | 1,623 | 1810 | -1.162148 | 30.143883 |
| Kashambya | 68 | 721 | 789 | -1.205614 | 29.935165 |
| Rwamucucu | 71 | 713 | 784 | -1.20972 | 30.06 |
| Hamurwa T/C | 116 | 582 | 698 | -1.206021 | 29.933555 |
| Muhanga T/C | 42 | 276 | 318 | -1.171253 | 30.124587 |
| Ruhija | 8 | 382 | 390 | -1.212469 | 29.949936 |
| Butanda | 24 | 403 | 427 | -1.32322 | 29.863589 |
| Katuna T/C | 304 | 271 | 575 | -1.428654 | 30.013897 |
| **Total** | **4,996** | **15,810** | **20,806** |  |  |

**Fig. 2:**
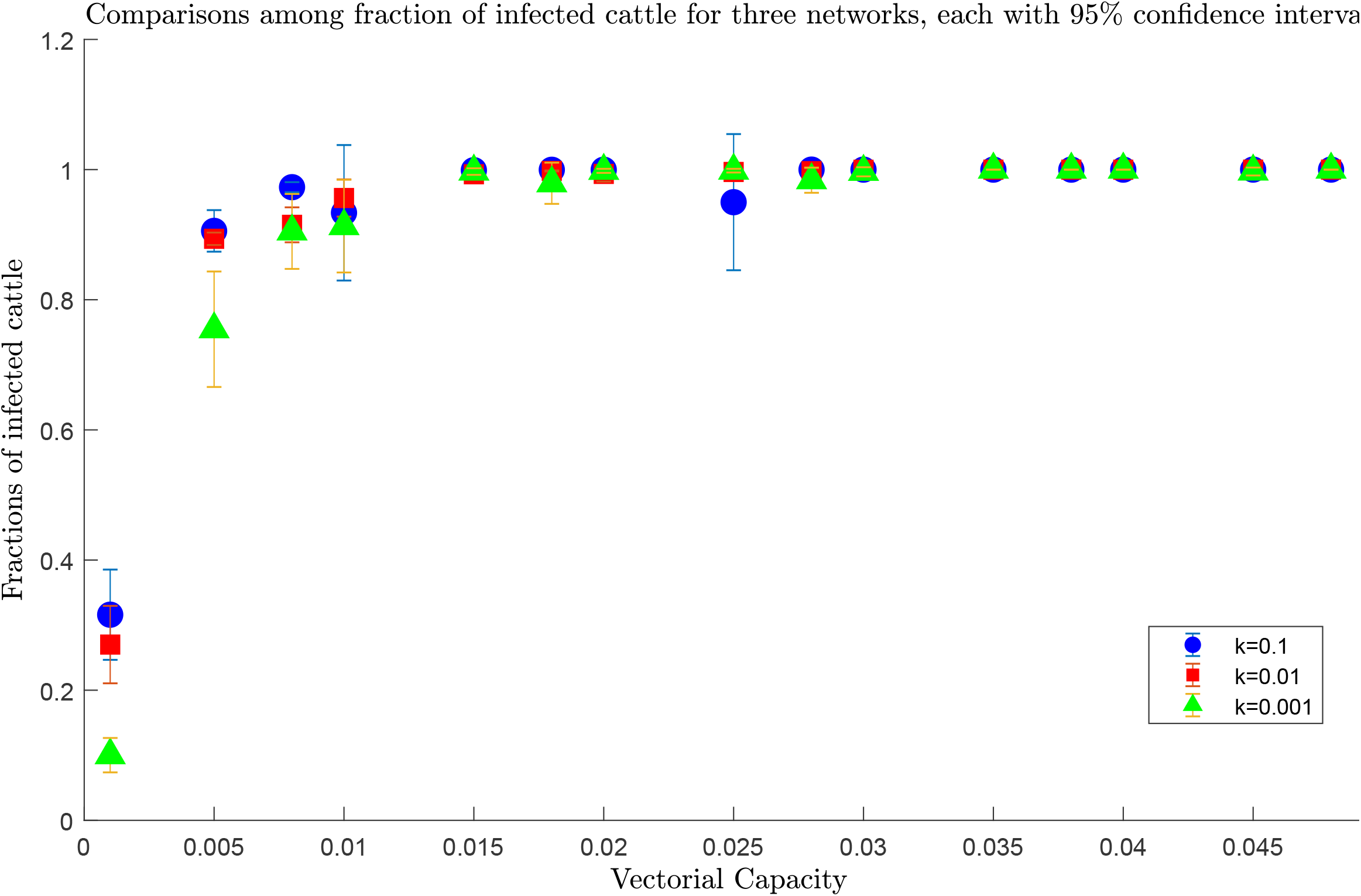
Location of each sub-county in Kabale District. Circles represent cattle farms in each sub-county and are color coded and scaled according to the total number of cattle in each location.

Some sub-counties were represented by data from a municipality (Kabale) or town council (Hamurwa, Muhanga, and Katuna) boundary (**Table 1**), and we extracted the longitude and latitude of the centroid of each sub-county, municipality, or town council boundary from Google maps to display in a GIS as shown in **Fig. 2**.

To capture the realism of the Uganda cattle movement and trading system in the model, we treated contact among cattle (not physical, rather implicit contact via mosquitoes) differently depending on geographic scale: cattle were assumed to move freely within each sub-county, while their movement was restricted between sub-counties. Within a sub-county we assumed each cow had equal probability to be connected to all other individual cows in that sub-county via mosquitoes because of their close relative proximity when compared to the Kabale District as a whole. This relationship among cows within sub-counties was best represented by an Erdos-Renyi network, where each node (i.e., cow) has equal probability of connectivity to any other node [15, 16].

Contact among cows from different sub-counties, on the other hand, called for a different kind of representation. Transmission of RVFV from one sub-county to another can happen via the movement of cattle for economic reasons, most commonly through cattle sales at local market places. Thus, contact among cows and thus the possibility of virus transfer from more proximal sub-counties was weighed higher than contact among cows and potential virus transfer from more distant sub-counties for this local trading system. We accomplished this weighting in the model with an exponential distance kernel, expressed as *e^-kd^*, where *k* is a constant which scales the probability of cows from different sub-counties to be in contact and has a unit *km^-1^*, and where *d* is distance between the origin and destination sub-locations.

We did not include the interactions of cows from different locations at market places in the transmission part of the model. Rather, we modeled the potential transmissions of RVFV that result from cows that move from their original location to another location, where they come in contact with others in that location. Therefore, an infected cow being moved to a new location can infect other cattle via local mosquitoes at the destination location; but we assumed no virus transmissions happen at the market place.

### Cattle Contact Network Visualization

We visualized the 20,806 cattle across the 19 sub-districts using the network visualization software Gephi [11], but scaling cattle population sizes across the network by a factor of 1/20 for clarity. It is important to note that scaling was only used for visualization and not model simulations which were performed with the full value of *N*. In **Fig. 3** the contact network visualization is shown where the name of the sub-counties are marked. The dense circular masses of varying size in **Fig. 3** represent the varying abundance of cattle at each sub-county similar to the scaled symbols shown in **Fig. 2**. Long black lines in **Fig. 3** connect some sub-counties, representing potential connections, and thus opportunities for mosquito-mediated transmission of RVFV between cows via cattle movements. The inset in **Fig. 3** expands a small portion of the contact network showing that the dense circular masses are actually made up of small black circles, each of which represents 20 cattle and correspond to the nodes shown in the representative contact network in **Fig. 1**. Likewise, the black lines among these nodes represent possible connections within and between sub-counties in the **Fig. 3** inset. Importantly, the particular arrangement of black lines (connectivity) in this visualization only shows one set of possible connections within and between sub-counties; modeling simulations consider many arrangements.

**Fig. 3:**
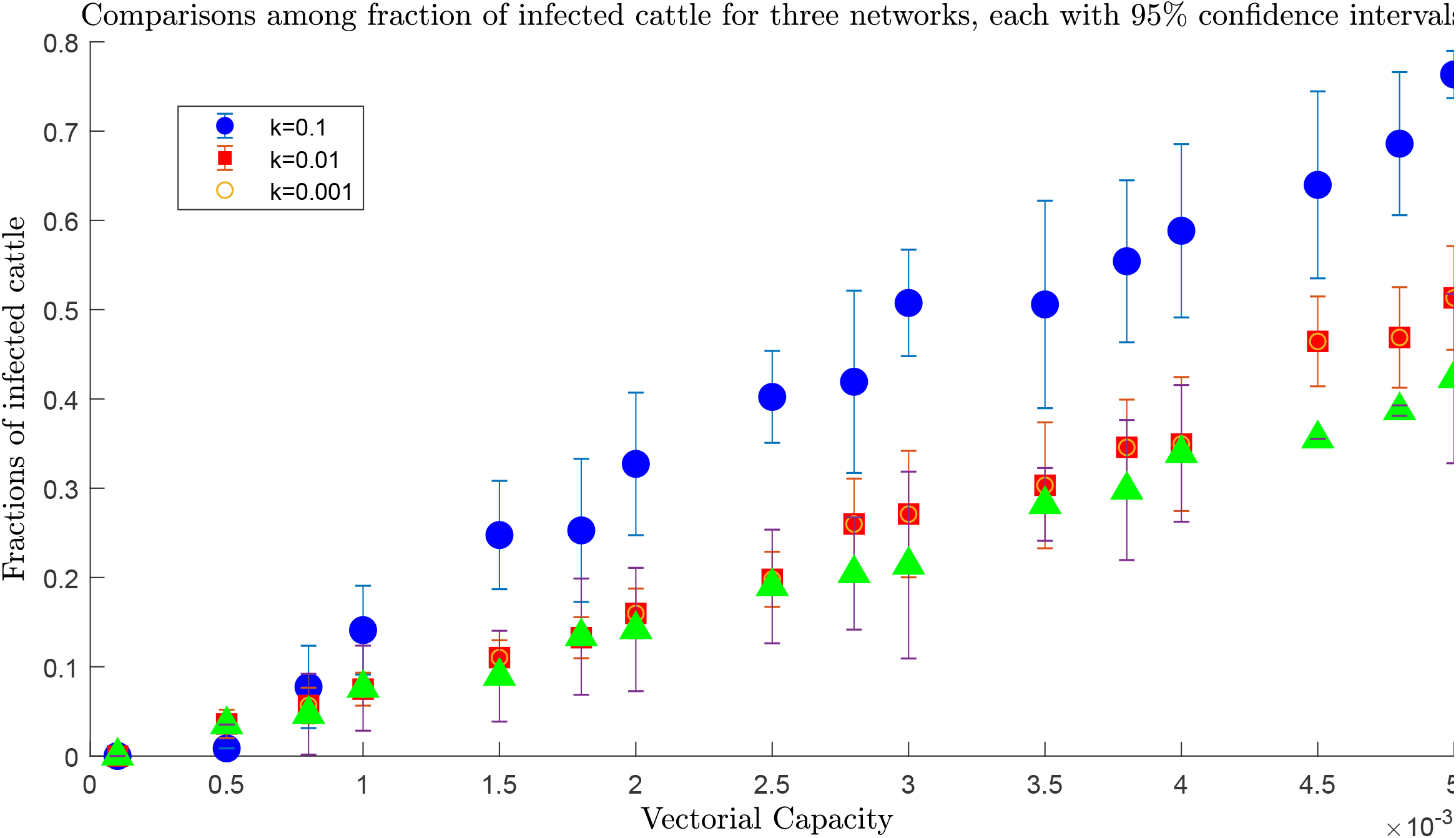
The overall structure of the network. Dense circular groupings of black dots represent different sub-counties. Inset shows close-up of two such groupings and one possible arrangement of links within and between them.

### RVFV Infection in Cows and Mosquitoes

We explicitly modeled the progression of RVFV infection of cattle with several key time components including: time for a newly infected individual cow to become viremic enough to infect a naïve mosquito feeding on it, time that the cow remains infectious, and time for the cow to become recovered. There is a link between two cows (i.e., a line between two nodes) if (a) virus transfer is possible between them (i.e., when one is infectious and the other susceptible), and (b) if they are in physical proximity; and virus transfer ultimately happens via infected local mosquito species competent for transmission of RVFV. Thus, links connecting nodes represent possibilities of RVFV transmission from an infected cow to a susceptible cow by a mosquito. The complexity and distance of connectivity from an infectious cow to susceptible cows across the contact network depends on human-mediated movement of cows among near or distant livestock operation locations (**Fig. 3**). The connectivity from this infectious cow to other susceptible cows within its original livestock operation location depends on the size of this original location.

Unlike the specific modeling of the progression of RVFV in the cattle, we only implicitly modeled RVFV progression in mosquitoes via a single parameter, transmission rate (*β*). Transmission rate is proportional to realized vectorial capacity of competent mosquito species likely to be present in the study area, and includes all key time components of RVFV infection in vector mosquitoes. Vectorial capacity is an aggregated measurement of the efficiency of vector-borne disease transmission which inherently considers climate and environmental factors: the Garett-Jones [12] equation for estimating vectorial capacity *C* is given as, which takes into account mosquito vector density proportional to host density (*m*), daily probability of host being fed upon (*a*), probability of daily survival of the vector (*p*), length of the virus extrinsic incubation period in days (*n*), and vector competence, or, the proportion of mosquitoes able to transmit RVFV (*b*) [9, 12].

In our model we estimated a composite index of vectorial capacity across indigenous mosquito species that takes into account survival and population density and can be estimated by real world data. If non-infected competent mosquito vectors are present and feed on newly arrived infectious cows, a fraction of these mosquitoes, proportional to vectorial capacity will become infected with RVFV, become infectious, and transmit the virus to immunologically naïve cows during subsequent blood feeding [9].

Our model does not account for heterogeneity of mosquito population distribution – for instance greater proximity of some livestock operations to immature habitat of RVFV mosquito vectors – and assumes all livestock operations are equally exposed to mosquitoes. On the other hand, the variation in mosquito transmission rates included in the model provides an aspect of heterogeneity in realized exposure to vector mosquitoes.

### Spread of RVFV Infection

In the network model, the infection can spread if a susceptible node (i.e., a susceptible cow) is in physical proximity with at least one infectious node. Specifically, one infectious cow (node 1) will be able to transmit RVFV to a susceptible cow (node 2) only if there are enough RVFV competent mosquitoes to first bite the infectious cow (node 1) then, after an appropriate period of time for the virus to disperse and replicate in the mosquito, bite a susceptible cow (node 2) [9].

As we have said above, the links between cattle in the network represent the feasibility of virus transfer via mosquitoes once cows are in physical proximity for a sufficient period of time – balanced by whether cows are naturally near each other within a sub-county or whether they come in contact due to trade-related cattle movement among sub-counties.

The node transition graph in **Fig. 1** represents the sequence of the progression of the RVFV infection in a cow (node) through four compartments, and is the core of the spread model: Once a susceptible (*S*) node is in physical proximity of an infectious node, virus transfer takes place with a rate *β* that is equal to the transmission rate, which is directly proportional to vectorial capacity, and moves the cow into the exposed (*E*) compartment. If a susceptible cow has *Y_i_* infectious neighbors, then the probability of the susceptible cow to receive virus transmission is *βY_i_*. Therefore, the total rate at which susceptible cows can become infected is proportional to the number of infectious cows in the neighborhood and the vectorial capacity of available mosquito vectors. The transition of the cow from the exposed compartment (*E*) to the infectious (*I*) compartment takes place at a rate δ, and represents the time the pathogen will take once it enters into the host body to replicate enough for the cow to become infectious – i.e., capable of infecting a naïve mosquito which may then infect a susceptible cow. Infectious cows are finally transferred to the recovered/removed compartment (*R*) with a rate *γ*. We did not distinguish whether RVFV-infected cows died or were recovered in this model; the endpoint in the simulation for an individual cow (node) was reached when it entered the *R* compartment.

Among these three transitions, the virus transfer from susceptible to exposed is called an edge-based transition because of the dependency on competent mosquito species, infectious cattle, and susceptible cattle. A transmission is possible only when there is a link between infectious and susceptible cattle. This link does not represent a physical contact between cattle, but rather the possibility of RVFV transfer via mosquito from infectious cattle to a susceptible one. The other two transitions, exposed to infectious and infectious to recovered, are called node-based transitions as they are dependent only on the host node.

After developing the individual-based SEIR network model for Kabale District, we carried out extensive simulations using Generalized Epidemic Modeling Framework (GEMF) that was developed by the Network Science and Engineering (NetSE) group at Kansas State University [4, 13]. In the SEIR model based on GEMF, infection processes were Poisson processes independent of each other. The node-level Markov process for node *i, i*= 1, 2,…*N*, is expressed as:

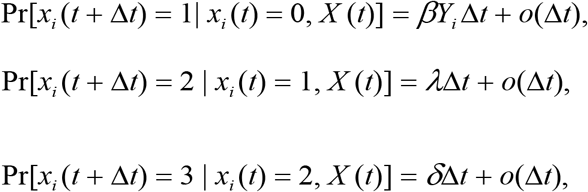

where *x_i_* = 0, 1, 2, or 3, which correspond to node *i* being in the susceptible, exposed, infectious, or recovered/removed state, respectively [9]. The value *X*(*t*) is the joint state of all nodes – the network state – at time *t*. In our model we used GEMF because it is individual-based which provides more accurate predictions than meta-population models [9] when we have real world data available. This individual-based GEMF model has also the ability to incorporate more complex network topologies when individual animal contact data are available.

### Cattle Network Scenarios

In the data set shown in **Table 1** two different kinds of cattle—*indigenous* and *exotic*— are accounted for in Kabale District, totaling 20,806 cows. There are cases in the literature indicating exotic cattle showed more susceptibility to RVF than indigenous cattle [14]. In Kenya, indigenous cattle exhibit fewer symptoms from RVFV infection and these cattle might also develop lower viremia which could significantly impact transfer of virus to mosquito vectors. However, we do not have specific information on relative susceptibility of indigenous compared to exotic cattle for the Kabale District in Uganda. This relative susceptibility could vary with breed as well as origin. Therefore, we assumed two different network scenarios to capture the relative susceptibility of exotic versus indigenous cattle breeds while performing simulations with GEMF: a *homogenous* network and a *heterogenous* network.

In the *homogenous* cattle contact network we considered that all cattle, indigenous or exotic, have the same susceptibility for RVFV. Therefore, we used the total number of cattle in each sub-county rather than differentiating them in two different categories.

In the *heterogeneous* cattle contact network we considered that exotic cattle are more susceptible to RVFV than indigenous cattle. Lacking the proper knowledge about the relative susceptibility, we assumed that if exotic cattle have a susceptibility *ζ*, then indigenous cattle have a susceptibility of *μζ*, where *μ* has a value between zero and one. For simulation purposes, we assumed *μ*=0.7. We can adjust the value of *μ* once real world data on relative susceptibility are available.

## Simulation Results and Discussion

We carried out extensive simulations under different conditions of cattle movement among sub-counties as well as different starting conditions to test the effectiveness of mosquito control and movement regulations, as well as the effects of cattle population diversity. We used the GEMF tool to run simulations with the networks described in *Materials and Methods*. For exponential connections between sub-counties, we used an exponential distance kernel *e^-kd^*, where *k* is the exponential constant and *d* is the distance between sub-counties. We assumed three different values of *k*, 0.001, 0.01, and 0.1, to reflect low, medium, and high movement probability, respectively. However, the network was valid for any value of *k*. We choose these values to explore the spectrum of varying movement probabilities in the network, and presented our simulations in three different sets, as follows.

For the first set, we presented simulation results for the three values of *k* as well as two ranges of the transmission rate *β*. Transmission rates incorporate mosquito abundance and weather factors such as humidity, temperature, and rainfall. Lacking sufficient data about cattle movement structure and mosquito vectorial capacity in Kabale District, we assumed these values according to Scoglio et al. [9] and Riad et al. [15] to explore a spectrum of cattle movement probability and mosquito abundance. Also lacking specific information about the value of *β*, we performed simulations for two different sets of *β* and two different network structures (*homogenous* and *heterogenous*) for each set of *β* and observed the variability in the fraction of infected cattle at the end of a period of 100 days, keeping weather factors constant and starting each simulation with only one infected cattle in the Kabale Municipality (**Fig. 2**).

For the second set of simulations, we showed the evolution of cattle in different compartments in the SEIR model by choosing a set of values of *β* (0.001, 0.005, 0.01, and 0.03) and starting with only one infected cattle in the Kabale Municipality (**Fig. 2**) for each simulation. The fraction of cattle in each compartment was plotted against time for these values of *β*. To reduce the number of simulations we choose medium cattle movement probability constant where *k*=0.01.

The third set of simulations consisted of investigating the effects of different starting conditions. We conducted simulations with different locations for the initial infected cattle, as well as single location and simultaneous multiple location epizootic outbreaks. We configured the network with values of *k*=0.01 and performed simulations for *β*=0.001, 0.005, 0.01, and 0.03 to reduce the number of simulation scenarios.

Therefore, we explored three different simulation sets, each consisting of a number of simulation scenarios, described as follows.

### Simulation Set I

In this set, simulations were initiated with a single infected cattle in the Kabale Municipality and conducted with three values of *k* (0.001, 0.01, and 0.1), two ranges of *β* (0001-0.005 or 0.001-0.048), and two network topologies *(homogeneous* and *heterogeneous*), producing four scenarios:

**Scenario 1:** *Homogenous* network and *β* range 0.0001-0.005
**Scenario 2:** *Homogenous* network and *β* range 0.001-0.048
**Scenario 3:** *Heterogeneous* network and *β* range 0.0001-0.005
**Scenario 4:** *Heterogeneous* network and *β* range 0.001-0.048

#### Scenario 1

The simulation for *β* ranging between 0.0001-0.005 (the lower range) is presented **Figs 4** for three different values of the exponential constant *k* and a *homogenous* network. We ran the simulation for 100 days and recorded the fraction of infected cattle for each value of *β*.

From **Fig. 4**, for *k*=0.01 and 0.001 we can see that after hundred days and for *β*=0.005 the infection reaches half the population. However, for *k*=0.1, after 100 days almost all of the cattle were infected. This is because the network was densely connected and it was easier for RVFV to be transmitted between individual animals than between sub-counties. This demonstrates that network structure plays a prominent role in RVFV spreading when the value of *β* is small. A value of *k*=0.1 means that extensive cattle movement between sub-counties will lead to infections in all cattle.

**Fig. 4:**
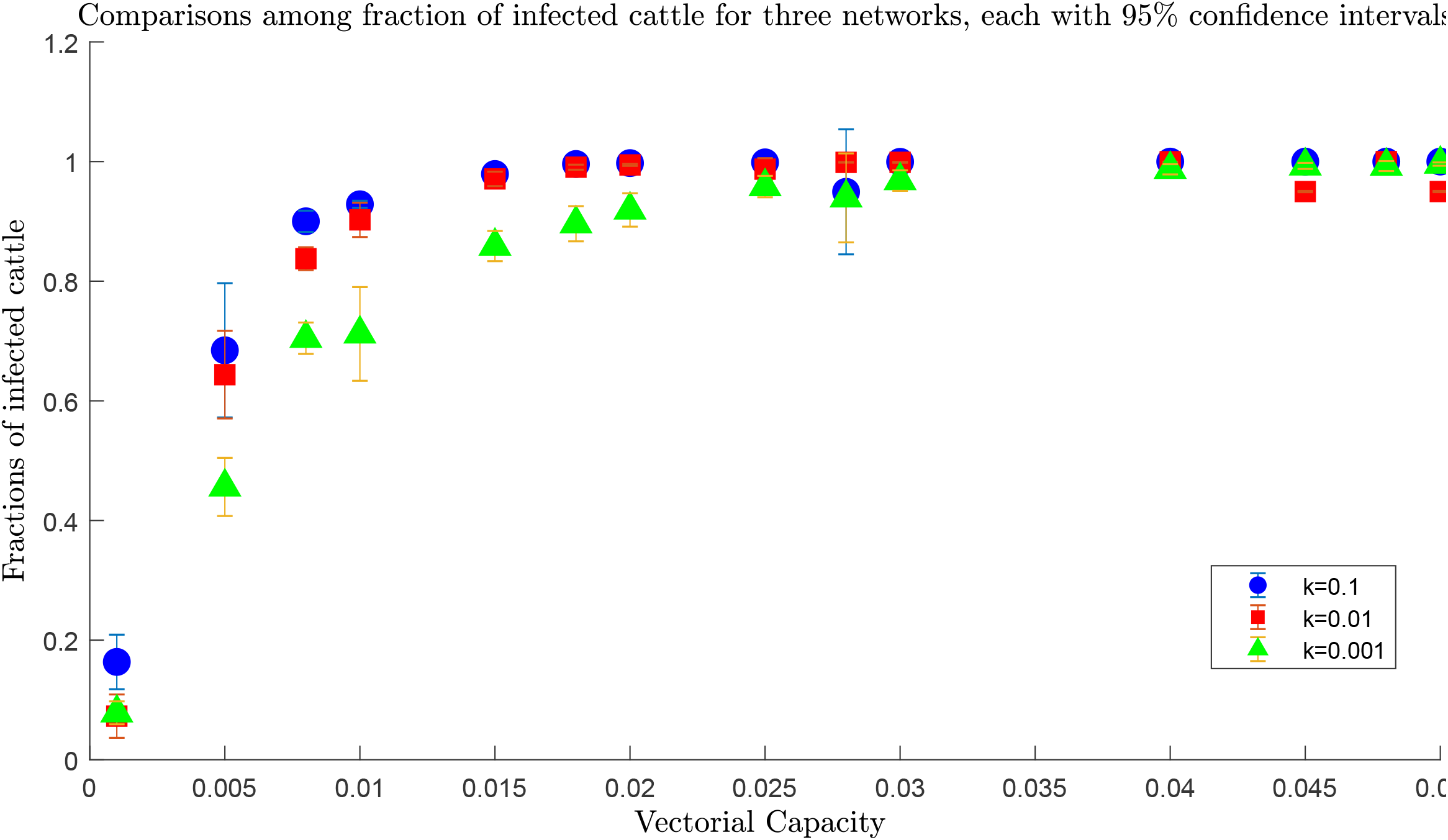
Comparisons among fractions of infected cattle for *homogeneous* network for three different values of *k* and lower range of *β*. Blue dots shows fraction of infected *k*=0.1, while red rectangles and green triangles showed fraction of infected for *k*=0.01 and 0.001 respectively. For the same value of transmission rate, we have always more infected cattle for greater value of *k* (0.1) than the smaller ones (0.01 and 0.001). Therefore, therefore increasing movement probability means more widespread epizootic, in particular for higher values of *β* such that the fraction of infected cattle at *β*=0,005 is ~0.399, ~0.537, and ~1 for k=0.001, 0.01, and 0.1, respectively.

#### Scenario 2

In the second set of simulations we used a *β* ranging from 0.001 to 0.048 (the upper range) for the *homogenous* network. Simulations results using these values are presented in **Fig. 5** for all three values of the parameter *k*.

**Fig. 5:**
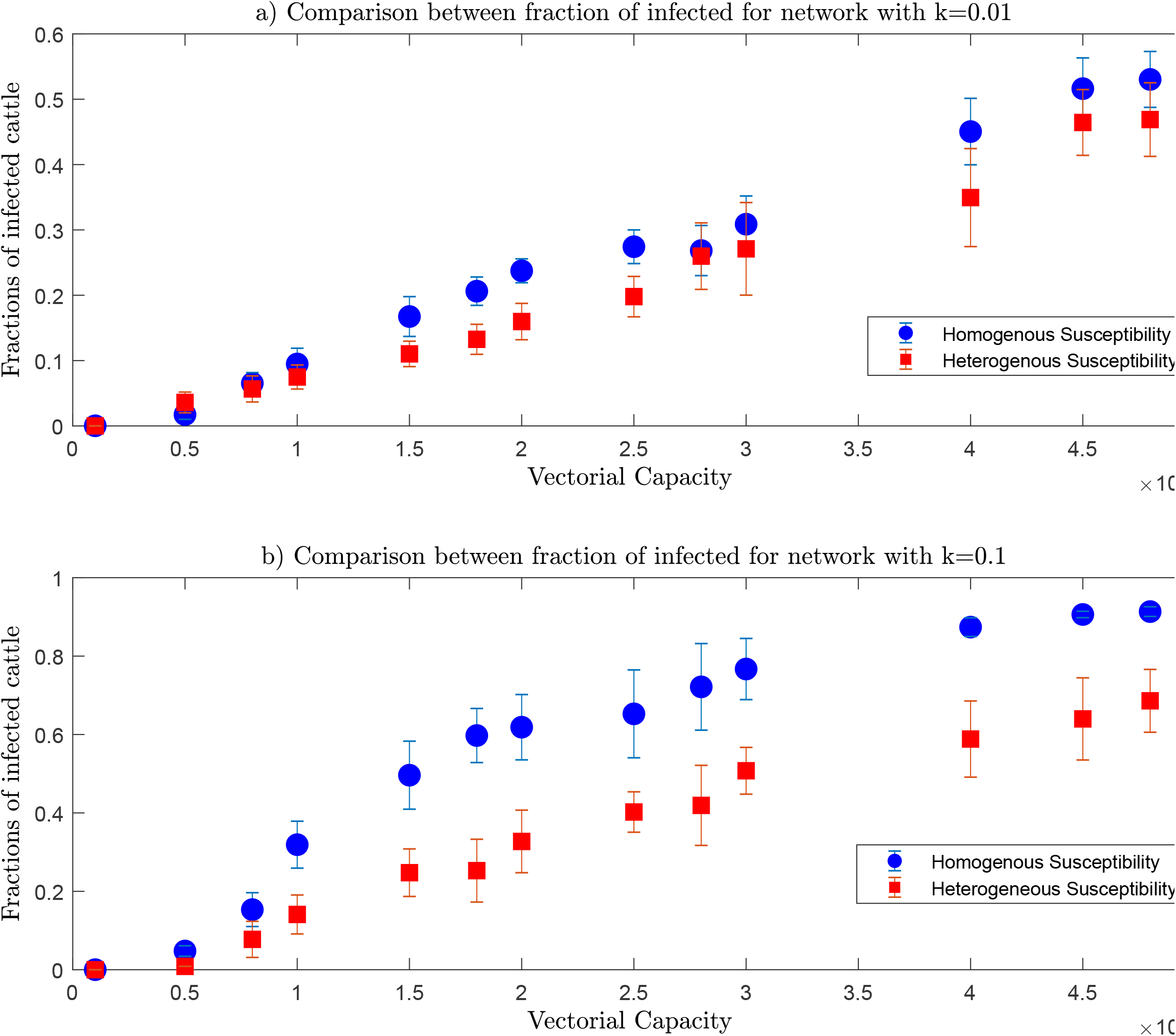
Comparisons among fractions of infected cattle for *homogeneous* network for three different values of *k* and upper range of *β*. Fractions of infected for all three values of *k* are almost overlapping, therefore, are not sensitive to the movement probability. They reach a value very close to 1, i.e., all 20,806 cattle became infected when transmission rate *β* reached 0.01 for the three networks. Therefore, fractions of infected cattle are also independent of the transmission rate *β* upper range.

From **Fig. 5** we can see that fraction of infected cattle reached 1 very quickly for this particular range of *β* for all three values of *k*. This is because we already have connections between sub-counties for all three networks. The whole cattle network becomes infected irrespective of the transmission rate (*β*) or cattle movement probability (*k*) the Therefore, we can say that for upper values of *β*, i.e., for higher abundance of mosquitoes and favorable weather conditions, the infection spread does not depend on the network structure and spreads throughout the whole network very quickly.

#### Scenario 3

In this scenario, we repeat the simulations for the lower range of *β* but for the *heterogeneous* network, and simulation results for three different values of *k* are presented in **Fig. 6**. There are increasing trends with the increase of *β* as well as *k* in the fractions infected cattle. For *k* 0.001 and 0.01, there is little difference; however, for *k*=0.1 the increase of the infected fraction is faster with increasing *β*. Therefore, if we want to reduce or check the spreading of RVFV, we need to reduce cattle movement. When we reduce cattle movement, only half the population becomes infected for the highest value of the transmission rate in the lower range. Therefore, reducing the cattle movement can reduce the number of infected cattle when there is a lower mosquito abundance and thus lower transmission rate.

**Fig. 6:**
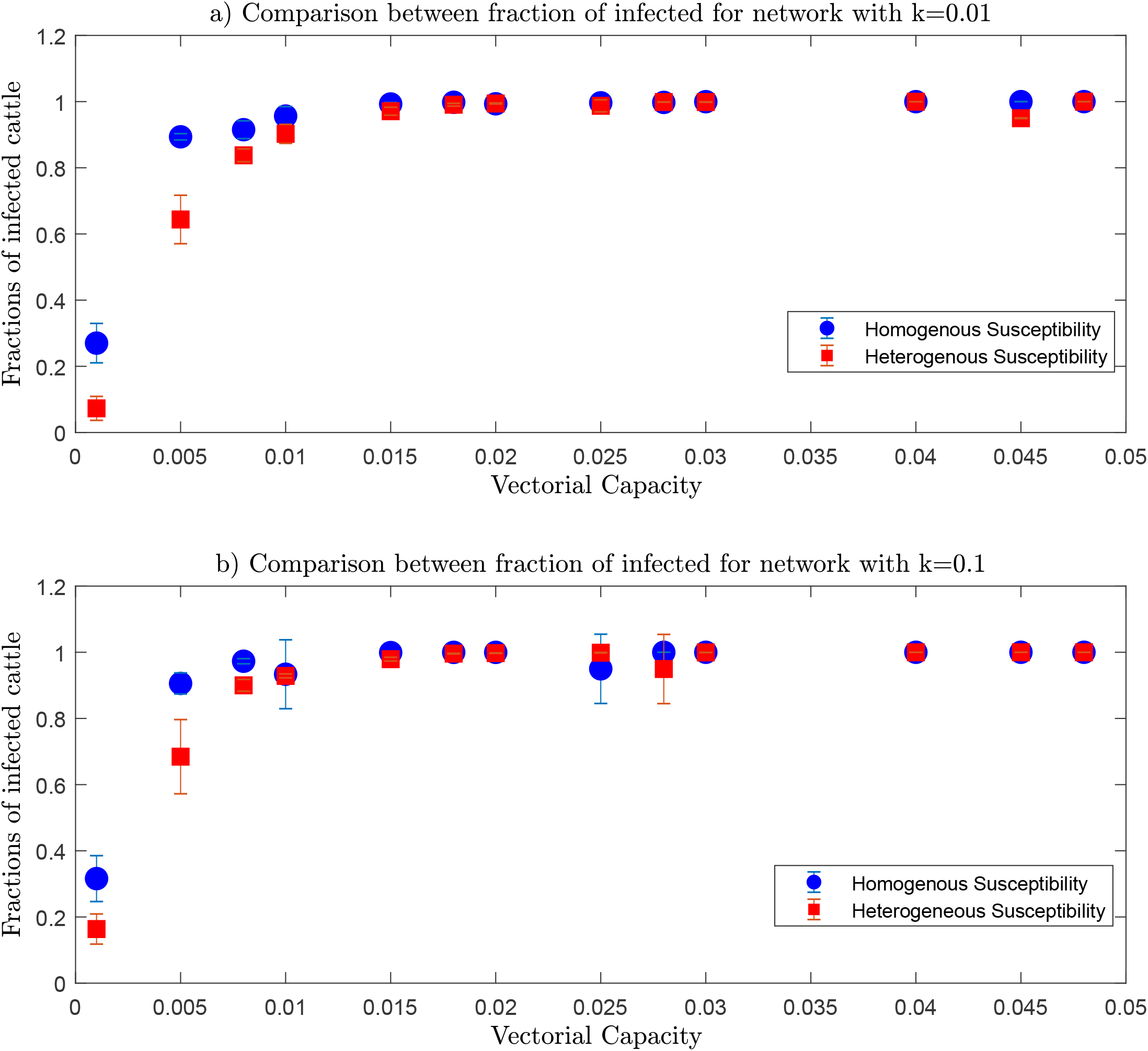
Comparisons among fractions of infected cattle for *heterogeneous* network for three different values of *k* and lower range of *β*. For *k*=0.001 and 0.01, maximum fraction of infected cattle is less than 0.5 for the highest value of transmission rate in the lower range, that means after the simulation period half of the cattle become infected. However for *k*=0.1, the infected cattle reaches up to 0.8. Therefore, we need to reduce the value of *k* i.e., cattle movement to reduce the fraction infected cattle

#### Scenario 4

Simulation results for the *heterogeneous* network and for the upper range of *β* are shown in **Fig. 7**. For all three values of *k*, the fraction of infected cattle reached 1 very quickly, near a *β* value of 0.02. After that, all 20,806 cattle become infected regardless of the values of *β* and *k*.

**Fig. 7:**
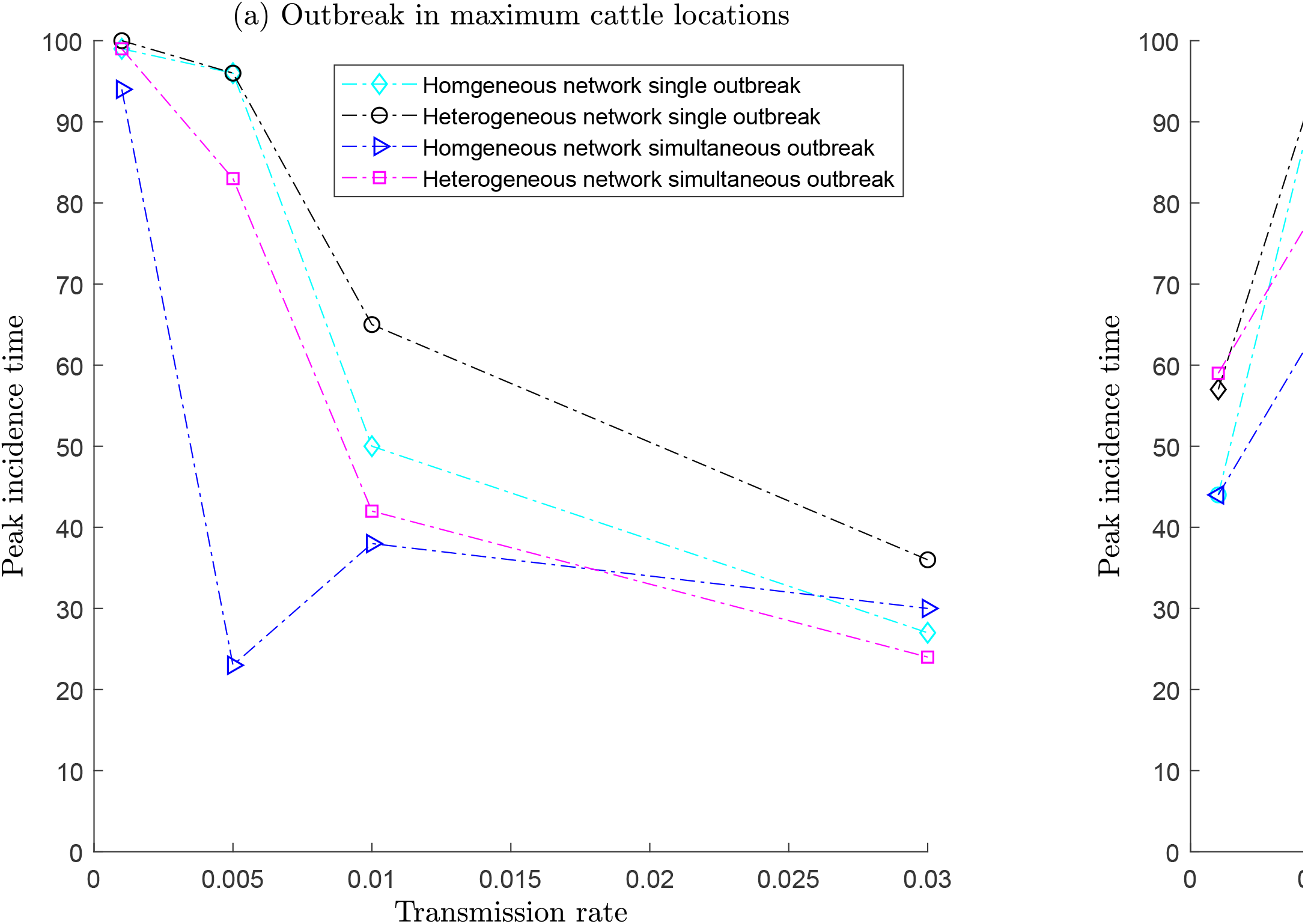
Comparisons among fractions of infected cattle for *heterogeneous* network for three different values of *k* and for upper range of *β*. The fractions of infected reaches towards one rapidly and when the value of transmission rate is 0.03, fraction of infected become one for three networks

The trend of the fraction of infected cattle for both *homogenous* and *heterogeneous* networks are similar in both lower and upper ranges of *β*. However, there are differences between fractions of infected cattle from *homogenous* compared to *heterogeneous* networks for same value of *k* and same range of transmission rate values. Comparisons between fractions of infected cattle for *homogenous* and *heterogeneous* networks are shown in **Figs 8** and **9**.

**Fig. 8** shows comparisons between fractions of infected for *homogeneous* and *heterogeneous* network for lower range of *β* values, and shows that the *homogenous* network has more infected cattle for the same values of *β* compared to the *heterogeneous* network.

**Fig. 8:**
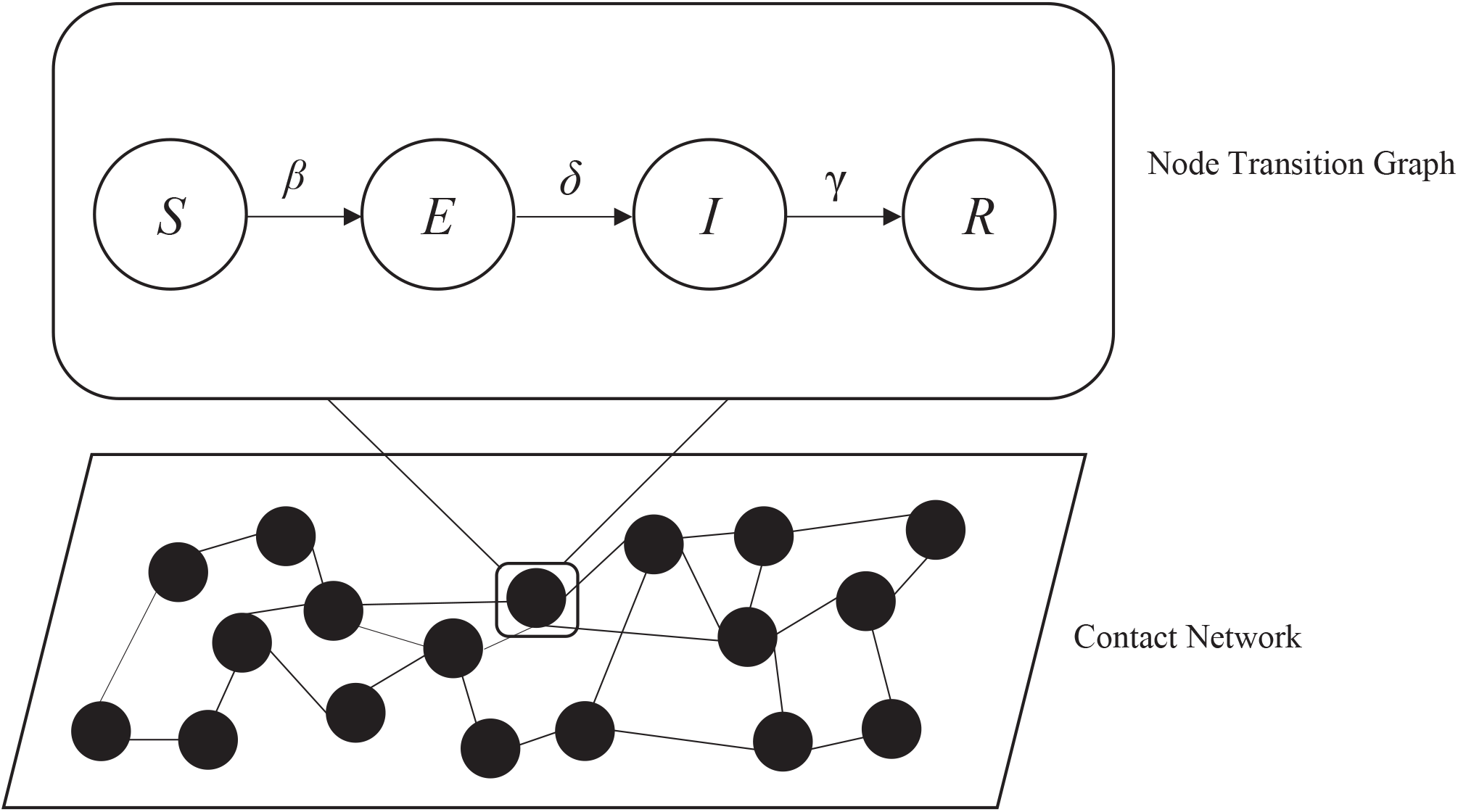
Comparisons among fractions of infected cattle for *heterogeneous* and *homogenous* networks for lower range of *β* and *a) k*=0.01 *and b) k*=0.1

In **Fig. 9** we can see similar outcomes, where more infected cattle result from the *homogenous* network. This can be attributed to the fact that, in the *heterogeneous* network, the indigenous cattle shows less susceptibility than the exotic cattle. Indigenous cattle species for being in that region for a very time, have developed certain immunity against RVF or other common diseases and they are more immune to RVFV than exotic cattle. Therefore, indigenous cattle are more likely not to get infected which results in a fewer number of infected cows than *homogenous* network where all animals are equally susceptible.

**Fig. 9:**
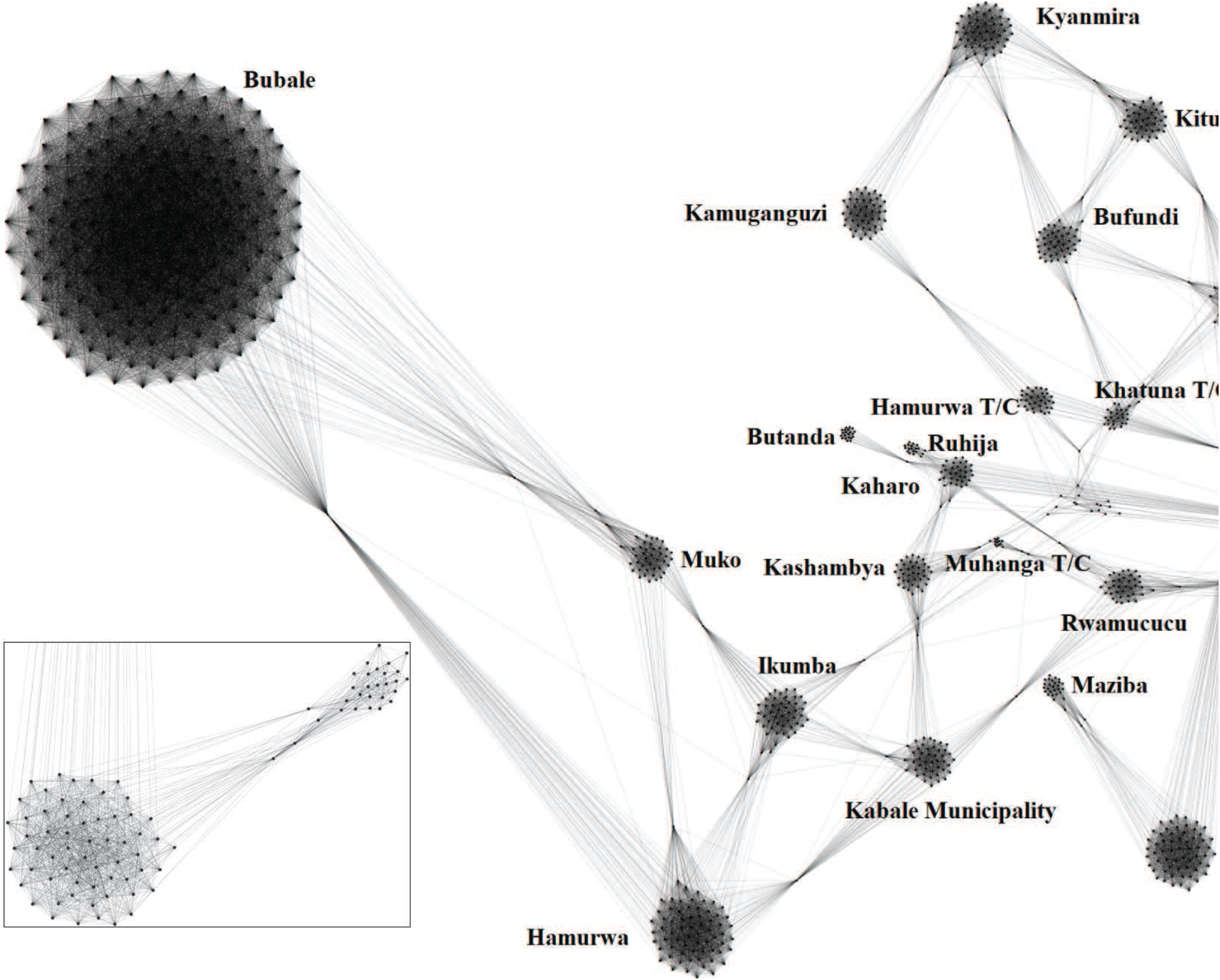
Comparisons among fractions of infected cattle for *heterogeneous* and *homogenous* networks for the upper range of *β* and a) *k*=0.01 *and b) k*=0.1

The comparisons between *homogenous* and *heterogeneous* networks show that reduced susceptibility of indigenous cattle to RVFV means a lower number of infected cattle during an RVFV epizootic. Therefore, greater proportions of indigenous cattle across sub-counties will reduce the numbers of infected cattle and thus a more contained epizootic.

In summary, simulations with lower transmission rates result in increasing fractions of infected cattle with increasing movement probability. However, for high transmission rates, the fraction reaches 1 quickly for *β*=0.01 and for *heterogeneous* network *β*=0.03 and there is little difference for increasing movement probabilities. From these observations we conclude that for low transmission rates (low mosquito abundance), restricted cattle movement will reduce the number of infected cattle. Higher transmission rates result in all cattle in the network becoming infected, regardless of cattle movement probability or mosquito abundance /transmission rate. These observations provide clear mitigation strategies to reduce the spread of RVFV. For a period of low mosquito abundance, cattle movement should be restricted to contain the epizootic to a minimum level; whereas, periods of high mosquito abundance (high transmission rates) require both mosquito control and cattle movement restriction. Comparisons between fractions of infected for *homogenous* versus *heterogeneous* networks suggest that diversity in the network results in fewer infected cattle for similar values of transmission rates and cattle movements.

### Simulation Set II

Simulations were conducted starting with a single infected cow in Kabale Municipality and simulations were conducted for a period of 100 d using both *homogeneous* and *heterogeneous* networks with *k*=0.01 and for *β*= 0.001, 0.005, 0.01, and 0.03 for each network, producing two scenarios:

**Scenario 1:** *Homogenous* network
**Scenario 2:** *Heterogeneous* network

We plotted the fraction of cattle in each compartment against time in Simulation Set II. For each scenario in this set, we assumed four different *β* to represent the entire range of transmission rates used for Simulation Set I. Instead of using different movement probability constants (*k*=0.001, 0.01, and 0.1) we choose *k*=0.01 for both *homogeneous* and *heterogeneous* networks.

#### Scenario 1

Simulation results for *homogenous* network and single infected cattle in Kabale Municipality for selected values of *β* are presented in **Fig. 10**. As we increase the value of *β* from 0.001 to 0.03, the fractions of recovered reach 1 very quickly. A very important point to be noted here is that fractions of recovered means these were the cattle who were infected in the first place. As we have not considered any disease induced mortality in the model, all infected cattle move to the recovered compartment. Therefore, the fraction of recovered cattle can be considered the cumulative fraction of infected for our specific model.

**Fig. 10:**
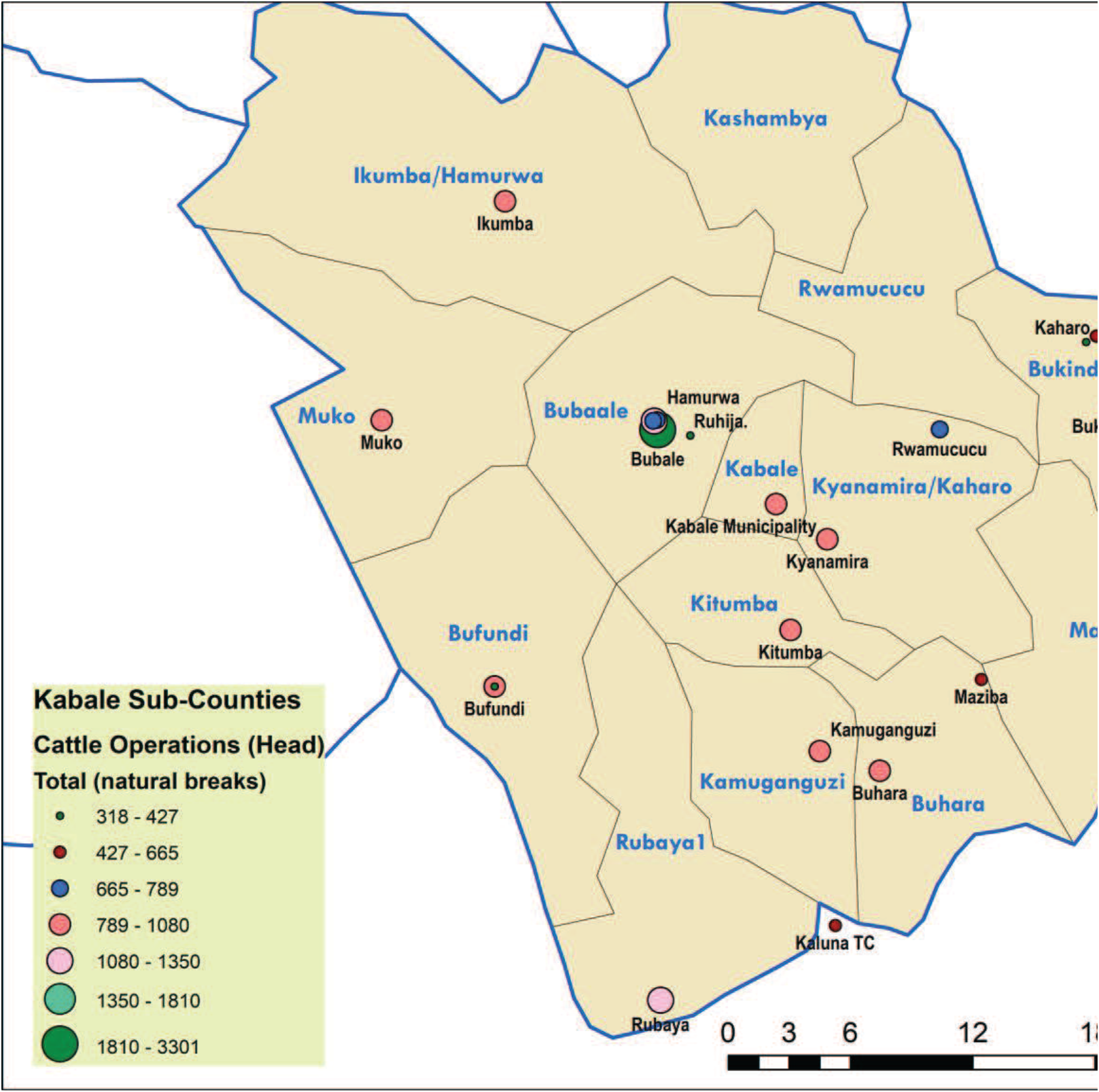
Fraction of cattle in each compartment with 95% confidence interval for *β*=0.001 (top left), 0.005 (top right), 0.01 (bottom left), and 0.03 (bottom right) and for *homogeneous* network. Increasing *β* shows an increasing trend in the overall fractions of recovered (cumulative fractions of infected)

**Table II:** Table showing maximum infected fractions of cattle, peak infection time, and rate at which that maximum is attained for *homogeneous* network.

| Transmission rate $\beta$ | Maximum infected fraction | Peak infection time | Rate |
| --- | --- | --- | --- |
| 0.001 | 0.0095 | 45 | 2.1268e-04 |
| 0.005 | 0.065 | 87 | 6.919e-04 |
| 0.01 | 0.0806 | 64 | 0.0013 |
| 0.03 | 0.1345 | 31 | 0.0043 |

#### Scenario 2

Simulation results for *heterogeneous* network with the initial condition of a single infected cattle in Kabale Municipality is presented in **Fig. 11**.

**Fig. 11:**
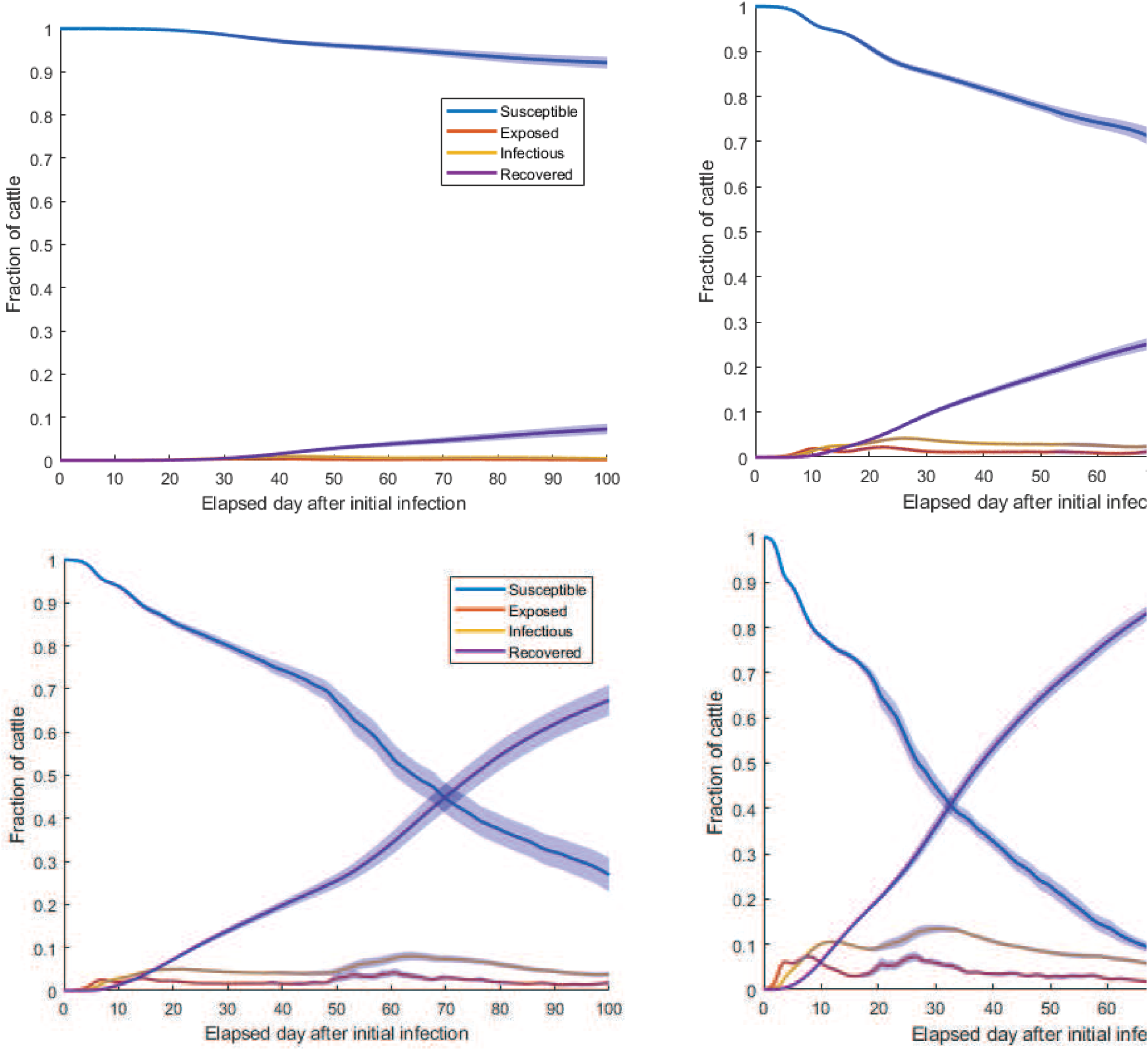
Fraction of cattle in each compartment with 95% confidence interval for *β*=0.001 (top left), 0.005 (top right), 0.01 (bottom left), and 0.03 (bottom right) and for *heterogeneous* network. Increasing *β* shows an increasing trend in the overall fractions of recovered (cumulative fractions of infected) which reaches to almost 1 for *β*=0.03.

**Table III:** Table showing maximum infected fractions of cattle, peak infection time, and rate at which that maximum is attained for heterogeneous network and a single infected cattle at Kabale municipality.

| Transmission rate $\beta$ | Maximum infected fraction | Peak infection time | Rate |
| --- | --- | --- | --- |
| 0.001 | 0.0056 | 60 | 9.333e-05 |
| 0.005 | 0.0365 | 100 | 3.6479e-04 |
| 0.01 | 0.0739 | 76 | 9.7690e-04 |
| 0.03 | 0.1181 | 43 | 0.0027 |

From **Table II** and **III**, it is evident that with the increase of *β*, the rate at which the fraction of infected reaches the maximum increases. However, there is an interesting trend that when the value of *β* is very small, i.e., *β*=0.001, the fraction of infected reaches the maximum faster for both *homogeneous* and *heterogeneous* networks than for a value of *β*=0.005. This can be attributed to the fact that, when the value of *β* is very small, the infection takes a long time to reach distant locations. Therefore, only cattle in the Kabale Municipality become infected within our simulation period of 100 d. When we increase *β*, infection reaches distant locations yet slower than the rate of infecting animals only in the initial location. However, when infection reaches distant locations, there are greater numbers of infected cattle in the network as a whole. This is evident from the maximum fraction of cattle, which is greater than the maximum fraction of infected cattle for *β*=0.001. Although the length of time taken for infections to reach maximum is greater for *β*=0.005 and 0.01, the rate always shows an increasing trend. Therefore, the increase in the transmission rate increases the severity of the epizootic. However, when *β* is sufficiently large (0.03), the time to reach maximum is less than time taken for *β*=0.001. Simulation results indicate that increase in the transmission rate expedites the spread of the epizootic in distant locations as well as the quantity of infected cattle. Therefore, mosquito control is crucial to contain the epizootic in the initial outbreak location while taking proper measures to care for infected cattle.

### Simulation Set III

These simulations were conducted with initial conditions other than starting simulation with a single infected cow in Kabale Municipality for both *homogeneous* and *heterogeneous* networks, as follows:

**Scenario 1:** Infection starts at 1 location (Bubale sub-county) with a maximum number of cattle
**Scenario 2:** Infection starts simultaneously at 3 locations (Bauble, Rubaya, and Hamurwa sub-counties) with a maximum number of cattle
**Scenario 3:** Infection starts at 1 location (Muhanga T/C) with a minimum number of cattle
**Scenario 4:** Infection starts simultaneously at 3 3 locations (Bukinda, Muhanga, and Ruhija sub-counties) with a minimum number of cattle

For detailed descriptions of simulation results from Simulation Set III, refer to the *Supporting Material*. We plotted the time to reach the maximum infection for each scenario with the transmission rate *β* in **Fig. 12**, which shows a brief summary of *Scenario 1* and *2* simulations when the initial RVF outbreak occurs in a single location (Bubale) or simultaneously at multiple locations (Bauble, Rubaya, and Hamurwa sub-counties), respectively.

**Fig. 12:**
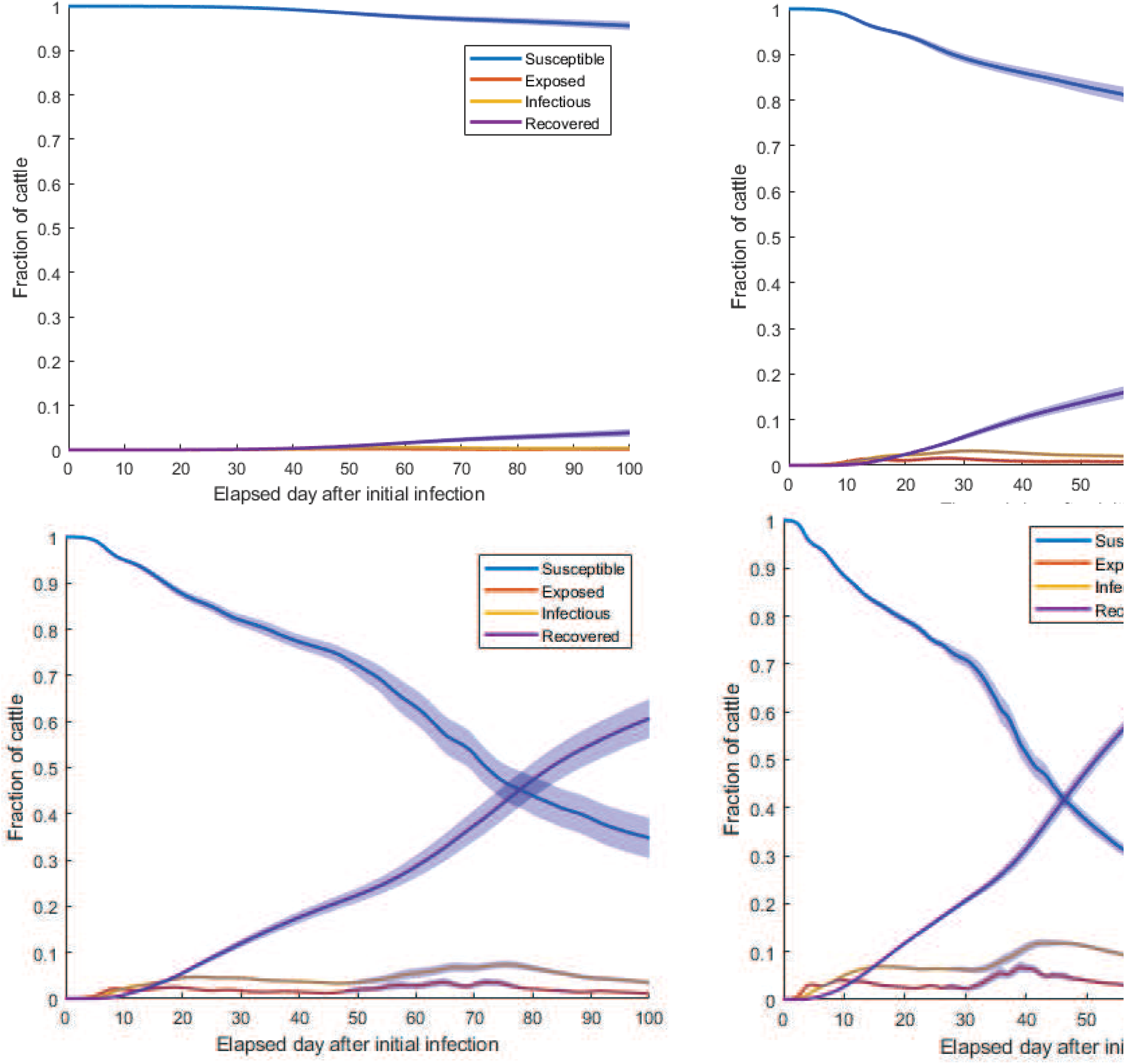
Peak infection time with transmission rate and for outbreaks starting in location/locations with (a) greater number of cattle and (b) fewer number of cattle.

Time taken to reach the maximum infection is smaller for simultaneous outbreaks regardless of the network structure than single-location outbreaks for similar values of the transmission rate *β* (**Fig. 12**). The spreading of infection through the network is always slower in the *heterogeneous* network for both single and simultaneous outbreaks. Infections spread slowly for the single-location outbreak in the network compare to the rate of spread in simultaneous outbreaks, which is reflected by the higher peak incidence time. For *β*=0.001, the peak infection time is close to 100 d for all simulations except simultaneous outbreaks in *homogeneous* networks (**Fig. 12**). This means that the peak has not been reached yet, and the number of infected cattle is still increasing. When we increase *β* to 0.01 (high mosquito abundance), the time to reach the peak reduces drastically for all of them.

An interesting trend emerges from **Fig. 12a:** when the value of transmission rate increases to 0.03, there is an increase in the time to reach maximum. This can be attributed to the fact that, when *β*=0.03, after our 100 d simulation period, all cattle were infected. Therefore, there are several time points when we have a peak in the fractions of infected animals at the same time (cattle recovered once they are infected, they stay in the infected compartment for a short period of time). The first peak infection can occur when infection spreads only in the initial location where the outbreak occurs, and the second peak can occur when infection spreads only to densely connected locations. Greater number of cattle location is densely connected with other location cattle in the network. Therefore, given enough mosquito abundance, the infections keep spreading to distant locations which result in an all-inclusive but a comparatively slower time to reach the peak for *β*=0.03. Here, a very important point to be noted that if we allow the simulation to run for a longer period of time, then we may observe a similar pattern of the peak infection time for all values of *β*. However, we limit our scope to 100 d simulation period in these simulations.

**Fig. 12b** represents peak incidence time when the RVF outbreak occurs in location/locations with fewer cattle than other locations. For lower mosquito abundance (*β*=0.001), infection does not reach distant locations, rather it is quickly confined to the initial location/locations, as is evident from smaller values of the peak infection time. However, with increase of *β*, the peak time returns to its regular pattern shown in **Fig. 12a**. We also don’t see any aberrant behavior for any selected scenario as seen in **Fig. 15(a)**, outbreaks being started in locations with sparse connections to other locations.

## Conclusions

Simulation results across multiple scenarios within these three simulation sets provide us important mitigation and intervention strategies. When a RVF outbreak occurs in a location with greater number of cattle, the infection spreads faster while infecting greater numbers of cattle than when an outbreak occurs in a location with fewer cattle. Therefore, the more the cattle in the initial outbreak location, the faster the spread and the more severe the epizootic. Simultaneous outbreaks in multiple locations will result in more severe and faster spreading of the epizootic than outbreak in a single location. That is evident from the values of the time to reach the maximum infection (**Fig. 12**) and rate at which the maximum infection is reached (refer to Tables in *Supplementary Material)*. Simulation results for different initial conditions and from both *homogeneous* and *heterogeneous* networks reveal similar patterns. Given the same initial conditions, the *heterogeneous* network is less susceptible to infection than the *homogeneous* network. This is evident from the rate at which infection reaches maximum fraction of infected, the value of the maximum infected fractions, and total cumulative fraction of infected.

We have used the cattle data for Kabale District and created a cattle movement network following an Erdos-Renyi topology inside each sub-county and exponential topology for inter sub-county movement. We have created two different networks— *homogenous* and *heterogeneous* – depending on the relative susceptibility of indigenous and exotic cattle. We used three different exponential parameters for each of the two networks to explore different spectra in inter sub-county movement probability. These networks were then used to run individual-based simulation using the GEMF tool developed in K-State NetSE group. From the simulation results, we saw that when the transmission rate is low, then the spread of RVFV throughout the network is dependent on inter sub-county movement probability. The more the probability, the more the fraction of infected cattle after the 100 d simulation period. This means that during periods of low mosquito vectorial capacity there must be more movement of cattle for a widespread epizootic. Therefore, during periods with reduced vectorial capacity, prohibition of inter sub-county cattle movement will eventually contain the epizootic in the initial location of the virus introduction. However, for upper values of transmission rate, i.e., upper mosquito vectorial competence, there is an increased likelihood of more extensive RVFV transmission that is less dependent the network structure (i.e., movement probability). Although, the risk of a widespread RVF epizootic is low in Uganda as demonstrated historically, when mosquito vectorial capacity is elevated it becomes critical to control/reduce mosquitoes to prevent widespread RVFV transmission. From the comparison of simulation results from *homogenous* and *heterogeneous* networks, we saw that for the same level of vectorial capacity and inter-subcounty movement probability, *heterogeneous* networks result in less infected cattle than *homogenous* ones because of the assumed reduced susceptibility of the indigenous cattle than exotic species. Therefore, the more indigenous cattle we have, the less will be the number of infected animals in the case of an RVF outbreak.

When infection starts in a location with more infected cattle then other locations, the epizootic spreads much faster than when infection starts in a location with fewer cattle. Simultaneous outbreaks in multiple locations cause significant increase in the rate to reach in the maximum infection spread. The rate increases with the increase of the transmission rate as well as the value of cattle movement probability. The location of the outbreak also plays a major role in the severity of an RVF epizootic. The simulation results from different initial starting locations showed that simultaneous outbreaks in different locations result in more cumulative infected cattle as well as faster rate of spreading compared to a single location initial outbreak. Population size in the initial location is also a crucial factor. The greater the population in the initial location, the faster the spread as well as the more the cumulative fraction of infected. Our simulation results quantitatively show how long the infection would take to reach maximum for different network structures as well as different starting conditions. Mitigation intervention can be devised from these simulation results as they provide us with the quantitative disease dynamics for epizootic spread. The longer the time to reach the maximum infection, the more time public and veterinary health personnel have to apply mitigation strategy before the infection becomes widespread. According to simulation results, applying insecticides to reduce mosquitoes will increase the spreading time of the infection and a greater opportunity to contain the epizootic by implementing mitigation strategies such as culling/removing infected cattle from contact with healthy animals via competent mosquito populations.

## Acknowledgements

We would like to thank the U.S. Department of Agriculture (USDA) Borlaug International Agricultural Science and Technology Fellowship Program for funding the fellowship, Center of Excellence for Emerging and Zoonotic Animal Diseases (CEEZAD) Kansas State University for supporting the fellowship program and College of Veterinary Medicine International Programs Kansas State University for supporting the fellowship program

## References

1. Tran A, Trevennec C, Lutwama J, Sserugga J, Gély M, Pittiglio C, Pinto J, Chevalier V. Development and assessment of a geographic knowledge-based model for mapping suitable areas for Rift Valley fever transmission in Eastern Africa. PLoS neglected tropical diseases. 2016 Sep 15;10(9):e0004999.

2. World Health Organization. Influenza (http://www.who.int/mediacentre/factsheets/fs211/en/).

3. Nanyingi MO, Munyua P, Kiama SG, Muchemi GM, Thumbi SM, Bitek AO, Bett B, Muriithi RM, Njenga MK. A systematic review of Rift Valley Fever epidemiology 1931–2014. Infection ecology & epidemiology. 2015 Jan 1;5(1):28024.

4. Sahneh FD, Scoglio C, Van Mieghem P. Generalized epidemic mean-field model for spreading processes over multilayer complex networks. IEEE/ACM Transactions on Networking. 2013 Oct;21(5):1609–20

5. Davies FG, Linthicum KJ, James AD. Rainfall and epizootic Rift Valley fever. Bulletin of the World Health Organization. 1985;63(5):941

6. Anyamba, A., J.-P. Chretien, J. Small, C. J. Tucker, P. Formenty, J. H. Richardson, S. C. Britch, D. C. Schnabel, R. L. Erickson and K. J. Linthicum (2009) Prediction of a Rift Valley fever outbreak. Proceedings National Academy of Sciences 106:955–959

7. Linthicum KJ, Britch SC, Anyamba A. Rift Valley fever: An emerging mosquito-borne disease. Annual Review of Entomology. 2016 Mar 11;61:395–415

8. Tuncer N, Gulbudak H, Cannataro VL, Martcheva M. Structural and practical identifiability issues of immunoepidemiological vector–host models with application to Rift Valley Fever. Bulletin of mathematical biology. 2016 Sep 1;78(9):1796–827.

9. Scoglio CM, Bosca C, Riad MH, Sahneh FD, Britch SC, Cohnstaedt LW, Linthicum KJ. Biologically Informed Individual-Based Network Model for Rift Valley Fever in the US and Evaluation of Mitigation Strategies. PloS one. 2016 Sep 23;11(9):e0162759

10. Uganda Bureau of Statistics (http://www.ubos.org/).

11. Bastian M, Heymann S, Jacomy M. Gephi: an open source software for exploring and manipulating networks. Icwsm. 2009 May 17;8:361–2.

12. Garrett-Jones C. The human blood index of malaria vectors in relation to epidemiological assessment. Bulletin of the World Health Organization. 1964;30(2):241.

13. Sahneh FD, Vajdi A, Shakeri H, Fan F, Scoglio C. GEMFsim: a stochastic simulator for the generalized epidemic modeling framework. Journal of Computational Science. 2017 Sep 1;22:36–44.

14. Gould EA, Higgs S. Impact of climate change and other factors on emerging arbovirus diseases. Transactions of the Royal Society of Tropical Medicine and Hygiene. 2009 Feb 1;103(2):109–21.

15. Riad MH, Scoglio CM, McVey DS, Cohnstaedt LW. An individual-level network model for a hypothetical outbreak of Japanese encephalitis in the USA. Stochastic Environmental Research and Risk Assessment. 2017 Feb 1;31(2):353–67.

16. Erdős P, Rényi A, Sós VT. On a problem of graph theory. Studia Sci. Math. Hungar. 1966;1(215):C235.

